# Similar Synaptic and Current Input Properties of Aromatase and Nonaromatase Neurons in the Zebra Finch Auditory Telencephalon

**DOI:** 10.64898/2026.07.16.738905

**Authors:** Basilio Furest Cataldo, Pamela Anderson, Luke Remage-Healey

**Affiliations:** Neuroscience and Behavior, Center for Neuroendocrine Studies, University of Massachusetts Amherst, 639 N. Pleasant St., Amherst, MA 01003, United States

**Keywords:** songbird, aromatase, electrophysiology, neuroestrogens, intrinsic excitability, post-synaptic currents

## Abstract

Estrogens, commonly known for their role in sexual development and maturation, support a wide variety of brain functions, including cognition, neuroprotection, and sensory processing. For example, in a songbird auditory forebrain region, *caudomedial nidopallium* (NCM), neuroestrogens are elevated in response to conspecific vocal and social stimuli; in turn, elevated neuroestrogens in NCM are known to rapidly enhance auditory processing independent of sex. Despite the pivotal role of neuroestrogens for brain function, it remains largely unknown whether aromatase (neuroestrogen-synthesizing) neurons differ in their cellular and physiological properties from non-aromatase-expressing neurons, as might be expected from specializations seen in other steroidogenic cell types (e.g., adrenal cells). Therefore, we systematically profiled aromatase and nonaromatase expressing NCM neurons to determine how they might differ in synaptic and current input properties, using *ex vivo* whole-cell electrophysiology followed by immunohistochemistry to reveal aromatase expression. Current-clamp recordings revealed no differences between aromatase and nonaromatase neurons, aside from a modest divergence in the time to reach peak membrane afterhyperpolarization. Similarly, voltage-clamp recordings of post-synaptic currents revealed no significant differences between the two cell types, suggesting that they share similar synaptic input density. Therefore, aromatase and nonaromatase NCM neurons share similar intrinsic membrane properties (e.g. excitability) and synaptic input dynamics in the higher songbird pallium. Our findings leave open the possibility that diverging computational and/or modulatory roles for aromatase vs. non-aromatase neurons could be due to projection- or neurochemical-specificity in their afferents and efferents.

**Significance Statement:** Neuroestrogens, locally synthesized in the brain by aromatase-expressing neurons, modulate neural circuit function across diverse brain regions, yet it remains unclear whether these cells represent a specialized neuronal population. We used whole-cell electrophysiology and post hoc identification of recorded neurons in the zebra finch auditory forebrain to compare the intrinsic and synaptic properties of aromatase and nonaromatase neurons. Confirming and extending previous findings, we show that aromatase-expressing neurons are not defined by unique electrophysiological properties. Instead, they appear to be functionally integrated within auditory circuits, supporting an alternative idea that neuroestrogen signaling acts through flexible neuromodulatory mechanisms rather than specialized neuronal properties. These findings do not exclude the possibility that aromatase neurons receive selective modulatory input from other brain regions.

## Introduction

Aromatase (estrogen synthase) is expressed throughout the vertebrate brain in a highly region-specific manner (Naftolin, Ryan, & Petro, 1972; Balthazart et al., 1996; Forlano et al., 2001; Stanić et al., 2014). Thus, aromatase expression, and the capacity to produce neuroestrogens centrally, has implications for brain function (Stocco, 2012) and behavior (Trainor, Kyomen, & Marler, 2006; Shaw, 2018; Takahashi et al., 2018; Shay, Vieira-Potter, & Rosenfeld, 2018). Locally-produced neuroestrogens can act on rapid timescales (seconds to minutes), distinguishing them from classical genomic steroid signaling, consistent with neuromodulators capable of dynamically shaping synaptic and intrinsic neuronal properties (Balthazart and Ball, 2006; Remage-Healey, 2012; Cornil et al., 2012; Srivastava et al., 2013; Scarpa et al., 2023).

Aromatase neurons are molecularly heterogeneous (e.g. Balthazart et al., 1991) and their distribution is remarkably conserved across vertebrates (Newman, 1999; Goodson, 2005; O’Connell and Hofmann, 2011; Spool, Bergan, & Remage-Healey, 2022). In addition, neuroestrogens are regulated outside the context of traditional sex- or social-specific behaviors, and we now understand roles for neuroestrogens in learning and memory (see review, Rosenfeld, Shay, & Vieira-Potter, 2018), emotion (e.g. Takahashi et al., 2018), motoric function (Azcoitia, Mendez, & Garcia-Segura, 2021), and neuroprotection (McCullough et al., 2003; Meltser et al., 2008). This widespread range of actions and contexts suggests a fundamental and evolutionarily ancient role for neuroestrogens signaling in social and behavioral circuit function.

Aromatase activity fluctuates in response to sensory input, behavioral state, and social context, producing dynamic changes in local neuroestrogen concentrations that can influence neuronal excitability on minute-by-minute timescales (Cornil et al., 2005; Remage-Healey et al., 2008; Cornil, Ball, & Balthazart, 2012; de Bournonville et al., 2012; Remage-Healey et al., 2012; Remage-Healey, Choleris, & Blathazart, 2018; Frick & Kim, 2018). Notably, aromatase is localized in both cell bodies and presynaptic terminals, enabling spatially precise, synapse-proximal estrogen signaling (Balthazart, 1993; Saldanha, Remage-Healey, & Schlinger, 2011; Remage-Healey et al., 2011). Consistent with this neuromodulatory framework, pharmacological inhibition of aromatase activity within discrete brain regions disrupts social and sensory-driven behaviors, underlying the functional importance of ongoing, local estrogen synthesis independent of circulating steroid levels (Cornil et al., 2005; Taziaux et al., 2007; de Bournonville et al., 2020; Gervais et al., 2019; Court, Balthazart, Ball, & Cornil, 2022; Cournoyer et al., 2026).

The *caudomedial nidopallium* (NCM) of the Zebra finch provides an accessible model for studying neuroestrogen function within a defined sensory circuit. NCM is a Wernicke-like higher-order auditory region (Bolhuis, Okanoya, & Scharff, 2010) and it contains a substantial population of aromatase-expressing neurons (Saldanha et al., 2000; Ikeda et al., 2017; Schlinger, 1997). This region is critical for auditory learning and song memory, exhibits stimulus-selective immediate-early gene induction and conspecific song recognition and discrimination (Mello, Vicario, & Clayton, 1992; Bell, Phan, & Vicario, 2015; Soyman & Vicario, 2019; Macedo-Lima and Remage-Healey, 2020). Importantly, aromatase activity in NCM is rapidly modulated by social and auditory stimuli, as local E2 levels increase during auditory and social contexts while disruption of aromatase impairs neural coding and behavior (Remage-Healey et al., 2010; Krentzel et al., 2020; Fernandez-Vargas et al., 2024). These findings establish NCM as a key site where neuroestrogen signaling is embedded within sensory processing.

Since aromatase neurons are both sources and potential targets of neuroestrogens (Spool, Bergan & Remage-Healey, 2022), one might expect them to exhibit specialized intrinsic and/or synaptic properties reflecting their neuromodulatory function. In murine medial amygdala, (ARO+) neurons are more abundant in males and they receive more projections from the vomeronasal organ, yet their electrophysiological membrane properties are not different from aromatase-negative neurons (Billing et al., 2020). Similarly, recent work using whole-cell current-clamp recordings demonstrated that ARO+/- neurons in NCM are indistinguishable in their intrinsic membrane properties (de Bournonville et al., 2021). The current interpretation, therefore, is that aromatase neurons are not a distinct cell type *per se* but are instead embedded within neural circuits amongst various neuronal identities, with facultative dynamics in both aromatase expression and activity.

Building on observations by Billing and colleagues (2020), we hypothesized that ARO+ neurons exhibit distinct synaptic input properties compared to ARO- neurons. If ARO+ neurons differ from their neighbors in synaptic current input amplitude and/or frequency this would advance our understanding of how aromatase neuronal firing state might be subject to electrochemical/synaptic control. To test this, we performed *ex vivo* current- and voltage-clamp recordings from NCM neurons, followed by aromatase immunohistochemical identification of recorded cells. We replicated previous current-clamp findings from de Bournonville and colleagues (2021) and further show that ARO+ neurons are indistinguishable from ARO− neurons across multiple measures of synaptic input.

## Methods

### Subjects

Juvenile (30 and 60 post-hatch days, phd) and adult (>90 phd) Zebra finches (*Taeniopygia Guttata*) were used in this study; age was a non-significant covariate across all analyses (*see Statistical Analysis* and **Supp. Materials 1**). Protocols were approved by the Institutional Animal Care and Use Committee at the University of Massachusetts, Amherst. All animals were bred and raised in our aviary.

### Slice extraction, preparation and whole-cell patch-clamp

*Ex vivo* slice recordings were used to determine the synaptic input profile and intrinsic membrane properties of NCM neurons, following previously published procedures (Spool et al., 2021; de Bournonville et al., 2021). Twelve ZFs were used in this study, and a final 35 neurons were included in all analyses. Following rapid decapitation, brains were extracted, blocked, immersed in ice-cold cutting solution (87 mM NaCl, 25 mM NaHCO3, 2.5 mM KCl, 1.25 mM NaH2PO4·H2O, 10 mM Glucose, 75 mM Sucrose, .5 mM CaCl2, 7mM MgCl2; 310 mOsm; 7.4pH when saturated with 95% O2 / 5% CO2), and saggitally sectioned at 250 μm thickness using a vibratome (VT1000S, Leica Biosystems Inc.). A total of 3 sections were used for recordings per hemisphere, to ensure all cuts were within the lateral boundary of NCM (∼1mm from midline; Dagostin et al., 2015; Mello & Clayton, 1994). Once sectioned, slices were immediately placed in warm aCSF external solution (130 mM NaCl, 2.5 mM KCl, 1.3 mM NaH2PO4·H2O, 26 mM NaHCO3, 10 mM Glucose, 1 mM MgCl2, 2 mM CaCl2). After 30 minutes of aCSF bath, slices were transferred to bubbled aCSF at room temperature. For electrophysiology, slices were placed in a recording chamber that was secured on a fixed stage microscope (Eclipse FN1, Nikon) equipped with a water-immersion objective (CFI Fluor; 40X; NA = 0.8; WD = 2.0 mm; Nikon). While the slices were continuously perfused with bubbled aCSF (carbogen), neurons were visualized with a CID-optic enabled charge-coupled camera (QIClick; QImaging). Borosilicate tubes were used to make glass pipettes via a two-stage vertical puller (PC-10, Narishige International USA). Just prior to recording, pipettes were back-filled with a mixture of ice-cold internal solution (100 mM K-gluconate, 20 mM KCl, .1 mM CaCl2, 5mM HEPES, 5 mM EGTA, 3 mM MgATP, .5mM NaGTP, 20 mM Na2-Phosphocreatine·4H2) and Neurobiotin (3mM), and the tip resistance for pipettes was between 4.5-6.5 MΩ. Cells were allowed to stabilize for a period of 3-5 minutes after achieving a whole-cell configuration. Both voltage- and current-clamp modes were used to acquire signals while the cells were maintained at the resting membrane potential. Current-clamp recordings consisted of recording voltage changes in response to hyperpolarizing and depolarizing current injections (500-1000 ms) at 10 or 20 pA steps; current-clamp recordings were sampled at 40kHz. Voltage-clamp signals were sampled at 20kHz and used to passively detect current changes while cells were held at -70mV. Experimental control and data acquisition was handled by multi-channel acquisition software (PATCHMASTER; HEKA Elektronik) and digitized via a patch-clamp amplifier (EPC 10 USB; HEKA Elektronik). All data was then exported for analysis in MATLAB and Python (*see Data analysis)*. For all cells, voltage-clamp recordings always came after current-clamp recordings, followed by a depolarization protocol (minimum 500, 5ms 450-700 pA current injections) to facilitate Neurobiotin transfer for post-recording visualization (IHC, *see Methods*). Following all recordings, the recording pipette was carefully removed, and brain slices were drop-fixed overnight in 4% paraformaldehyde. The fixed tissue was then moved to a cryoprotectant solution and kept at -20 until used for immunolabeling.

### Immunohistochemistry

Immunohistochemistry was used to visualize the Neurobiotin-filled recorded neurons and to determine cell type (**Fig. 1**): aromatase-positive (ARO+) or aromatase-negative (ARO-). Cryoprotectant was first washed off with a series of 5 5-min PB rinses, then the tissue was permeabilized across 3 10-min .3% PBT (PB + Triton-X), followed by a 1-hour blocking period in a 10% normal goal serum in .3% PBT solution. Tissue was then incubated for 48 hours with a polyclonal rabbit anti-aromatase primary antibody (a gift from Dr. Colin Saldanha, American University; 1:2000) in a 10% normal goal serum in .3% PBT solution; the antibody has been previously validated in ZF tissue (Saldanha et al., 2000; de Bournonville, 2021). Next, after rinsing off the primary antibody solution across 3 10-min .1% PBT washes, the tissue was incubated with a solution of .1% PBT with Alexa-594-conjugated goat anti-rabbit antibody (ThermoFisher Scientific Inc., MA; 1:200) and dylight 488 streptavidin (Vector Laboratories SA-5488; 1/200) for 1 hour at room temperature. Lastly, secondary antibodies were washed off if .1% PBT, mounted onto Superfrost plus slides (Fisher Cat No. 22-037-246) and coverslipped using Antifade mounting medium with DAPI (Vectashield, H-1200-10).

**Figure 1.**
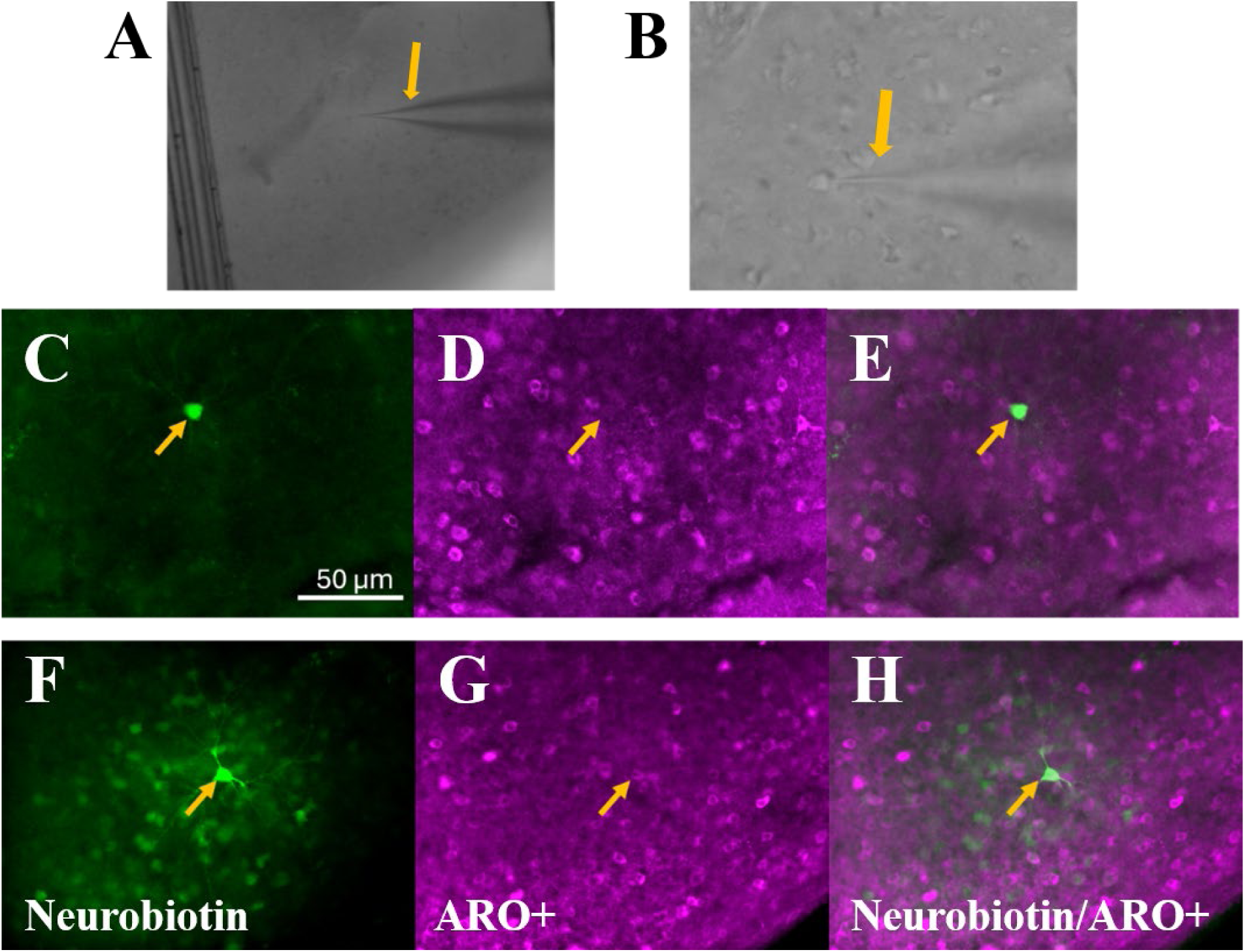
*Ex vivo* electrophysiology and histology. Example of patch electrophysiology with visible pipette (yellow arrow) at 10x (A) and 40x (B) magnification. Patched cells and their cell type (aromatase-expressing or not) was achieved by filling recording cells with Neurobiotin followed by immunohistochemistry. Cells were first visualized based on Neurobiotin expression (C and F), aromatase expression was revealed for every cell (D and G), and patched cells were determined to be aromatase positive if these signals colocalized (E and H). The first row (C-E) displays a patched cell that was ARO- and the second row (F-H) displays a patched cell that was ARO+.

Sections were imaged using a confocal microscope (NA1, Nikon, Tokyo, Japan) and an all-in-one fluorescence microscope (Keyence, BZ-X810). Visualization of Alexa 488 revealed which cell was recorded and Alexa 594 revealed which neurons expressed aromatase. Colocalization was determined manually while imaging tissue and confirmed *post-hoc* in FIJI (Schindelin et al., 2012). Aromatase expression was further confirmed by computing a ratio between the aromatase expression intensity of the target cell (by creating an ROI in the aromatase 594 channel that coincides where the biotin-filled cell is on the image) and the average intensity of the image on the same channel. Mood’s median test confirmed that this ratio was significantly different between cells labeled as aromatase-negative (median = .877, SD = .13) and aromatase-positive (median = .99, SD = .04), *χ*^2^(1) = 17.35, p < .001.

### Data Analysis: Current-clamp

Data files obtained from HEKA software were first fed through a MATLAB routine (https://github.com/ChristianKeine/HEKA_Patchmaster_Importer) to extract the data into an accessible data structure, and then the structure was read into Python for all analyses via custom-scripts (*available on GitHub*). Voltage-Current (VI) relationships were described by plotting changes in current against the corresponding changes in voltage. Input resistance (R_i_) was computed as the slope of this relationship by restricting the range to hyperpolarizing current steps. Rheobase was defined as the minimum amount of required current to elicit one action potential. Frequency was defined as the number of detected action potentials during the current injection, divided by the duration of the injection. The resting membrane potential was derived by finding the median membrane potential in the 3 seconds before the current injection at rheobase. The AP threshold was determined by smoothing the rheobase trace with a Gaussian filter, then finding the maximum value of the second derivative. The AP peak at rheobase was found at the first point at which the trace reached its max value. The afterhyperpolarization (AHP) of the first rheobase spike was picked out manually by finding the time and voltage of the lowest point after the peak. The width of the action potential was calculated by finding the midpoint between the threshold and the peak and then finding the time at which that voltage occurs between the peak and the afterhyperpolarization. These two values were then subtracted from each other for the action potential half-width. Lastly, neurons phenotype to be phasic if they fire a maximum of 3 action potentials across current injections, and tonic if they fired >3 action potentials across current injections (**Fig. 2**). Group comparisons between ARO+ and ARO- were conducted while assuming unequal variances via Welch t-test; Hedge’s g was used for effect size computation with un-pooled standard deviation. To ensure robustness to heteroscedasticity and unequal sample sizes across groups, all factorial analyses (cell type vs cell phenotype) were conducted via two-way robust ANOVAs with 20% trimmed means and robust estimators of variance (*t2way* in WRS2 R-package).

**Figure 2.**
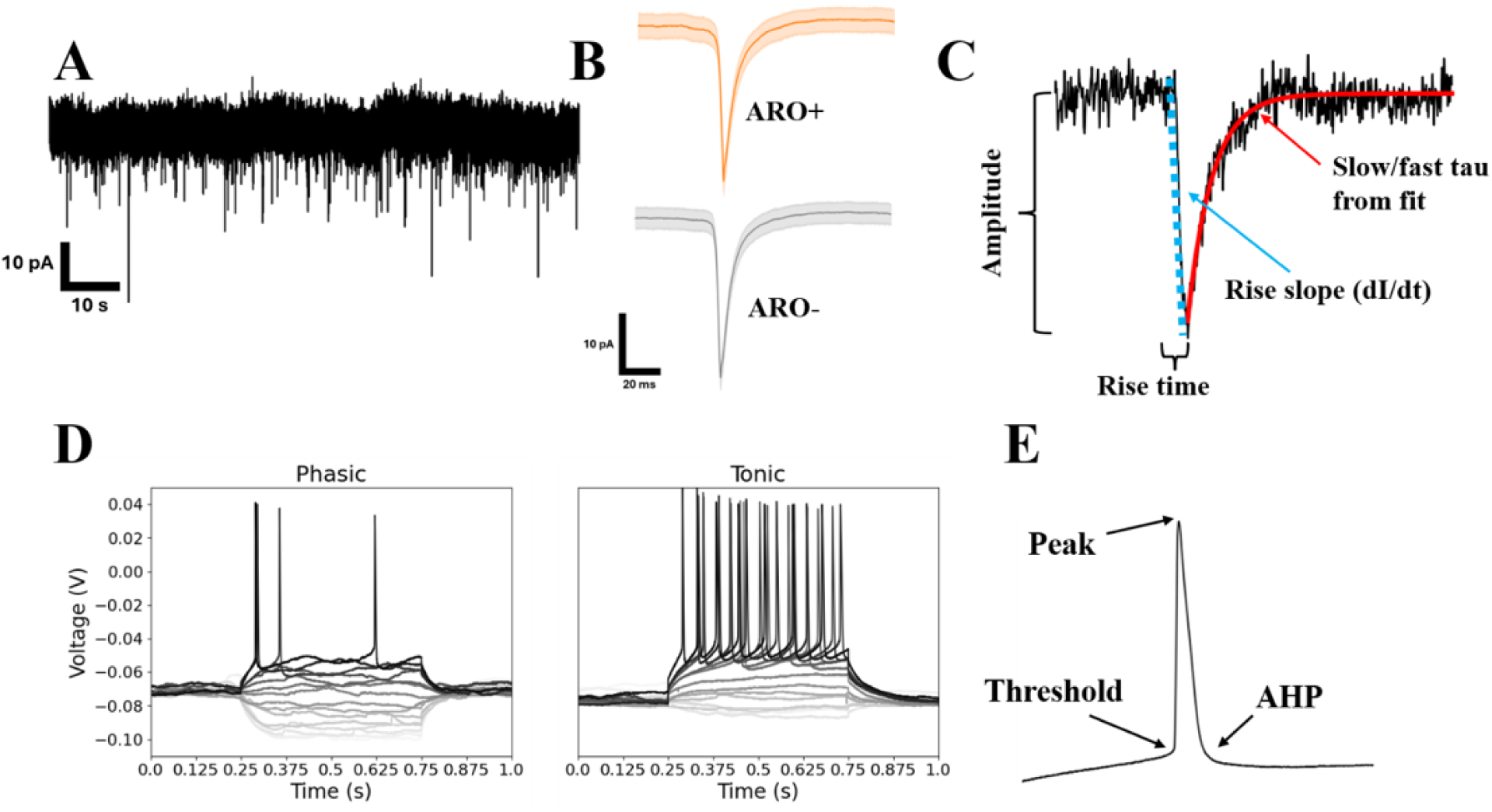
Voltage- and current-clamp recordings and measurements. A) Voltage-clamp trace over a 5-minute period of passive recording. B) Averaged PSC waveforms recorded from ARO+ (top) and ARO- (bottom) cells. C) Definition of PSC measurements. Amplitude reflects the difference between peak voltage and the median voltage of the preceding baseline. Rise time was defined as the period it takes from the base to the peak of the current event. Rise slope was defined as the slope from the base of the PSC to its peak (dI/dt). Lastly, slow and fast tau were derived, for each individual PSC, as parameters in a double-exponential fit. D) Example current-clamp recordings, different color denotes a different current injection at a step of 10pA. Phasic cells (left) were defined as cells that fired fewer than 3 action potentials, whereas cells were classified as tonic (right) if they fired more than 3 action potentials when driven by current injections. E) Close up of an action potential elicited during a current-clamp recording. Arrows denote key landmarks used to derive different measurements of action potential dynamics.

### Data Analysis: Voltage-clamp

Voltage-clamp analysis focused on inward post-synaptic currents (PSCs), which were detected with a recently published, GUI-driven python package (PatchView; Hu & Jiang, 2022). The software initially uses a deconvolution filter and a threshold to detect PSCs, which are then manually curated, and the data (PSC waveforms and timestamps) was exported to a custom python script. PSC amplitude was computed as the difference between the peak-voltage and the median-current during the PSC-preceding baseline. Rise time was derived as the time between the PSC peak and the moment at which it reaches the baseline. Rise slope was defined as the linear gradient from the base of the peak to the peak itself. Lastly, fast and slow tau were computed for the waveforms (**Fig. 2**). A differential evolution algorithm (derived by Storn & Price, 1997, and deployed in the SciPy library by Virtanen et al., 2020) with a sum-of-square error loss-minimizing function was used to fit a biexponential curve (**Equation 1**) onto each PSC separately. Adj-R^2^ of the fit offered a rough approximation of fit and data quality, and all analyses were carried out with PSCs that exhibited adj-R^2^ that were higher than the median, ∼57%. For each PSC, fast (first time component) tau and slow (second time component) tau were derived as the time coefficients of the double-exponential function. Lastly, peak frequency was derived as the sum of detected peaks divided by the recording length (in seconds). Graphical displays were made in Python and statistical tests were conducted using standard statistical packages in Python. Welch t-tests at an alpha of .05 were used to compare ARO+ vs ARO-neurons across the range of metrics (e.g. amplitude) that were first averaged per neuron, such that equal variances were not assumed between groups; Hedge’s g was used for effect size computation with unpooled standard deviation. General Linear Mixed Models (**Supp. Materials**) were employed to detect whether group differences arose while considering all possible PSCs from each neuron (i.e., not averaging PSC features), and to determine if finch age, as a covariate, contributed to a given metric. Ninety-five percent confidence intervals for each metric, based on median group differences using the Hodges-Lehmann estimator (Hodges & Lehmann, 1963), were developed to visualize their estimate differences.

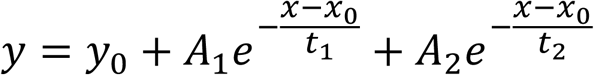

**Equation 1**. Biexponential fitting function for PSCs. *t*_1_ and *t*_2_ values were used as *τ_slow_* (slow tau) and *τ_fast_* (fast tau) respectively.

## Results

### Current-clamp recordings confirm similarity between ARO+ and ARO- neurons in intrinsic properties

Individual neurons’ intrinsic membrane properties, measured via *ex vivo* current-clamp electrophysiology, were compared based on whether the cell was ARO+ or ARO- (**Fig 3A**) and as a replication of previous work (de Bournonville et al., 2021). First, only one metric related to the temporal dynamics of these properties was observed to be significant: ARO+ (*x̃* = .161, SD =.06) was found to exhibit significantly, albeit modestly, higher AHP time relative to ARO- (*x̃* = .111, SD =.059), *t*(21.847) = 2.206, *p*. 038, *g* = .812. However, Time of Threshold was not different between ARO+ (*x̃* = .117, SD =.046) and ARO- (*x̃* = .085, SD =.058), *t*(25.078) = 1.654, *p* = .13, *g*. 596, nor was Peak Time between ARO+ (*x̃* = .119, SD =.046) and ARO- (*x̃* = .086, SD =.058), *t*(25.092) = 1.653, *p* = .131, *g* = .596. In this parametric approach, none of the voltage-based measurements—RMP, Voltage of Threshold, AP width, Voltage at AHP, Voltage of Peak—were significantly different (**Table 1**). Median-difference 95% confidence intervals based on the Hodges-Lehmann estimator for each metric can be seen in **Figure 3B**. To investigate whether cell phenotype (i.e. tonic vs phasic), cell type (ARO+ vs ARO-), and their interaction significantly explained a given current-clamp measurement, each metric was modeled via multi-factor HC3-controlled OLS. A two-way robust ANOVA with 20% trimmed means confirmed (as seen in previous analysis) that cell type is a significant factor for AHP Time (Q = 11.611, p=.018). Similarly, robust ANOVAs were conducted for each metric separately, with cell phenotype and type as factors in addition to their interaction. Cell phenotype was detected as a significant factor for both Threshold Voltage (Q = 6.921, p = .027) and AHP Voltage (Q = 10.32, p = .006). No significant interactions were observed. All results are tabulated in **Table 2** and visualized in **Figure 4**.

**Figure 3.**
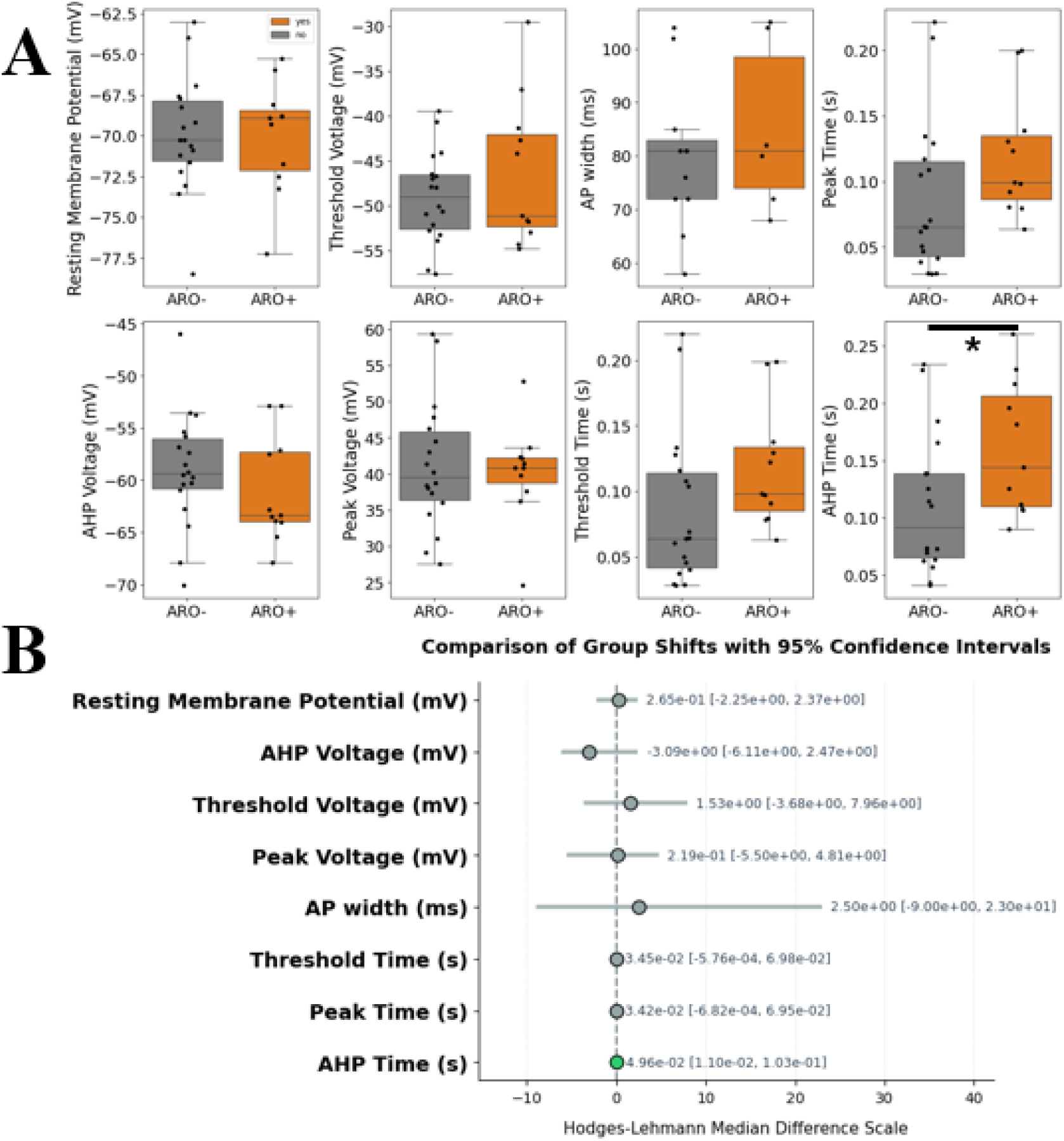
Comparison of voltage-clamp-derived PSC measurements between histologically-confirmed ARO+ (orange) and ARO- (grey) cells. A) While ARO+ cells displayed a modest yet significantly higher AHP time relative to ARO- cells, there were otherwise no other significant differences between the two cell types. Each datapoint reflects a cell. B) A visualization of the compared differences in terms of 95% confidence intervals. * reflects a significant difference at an alpha of .05.

**Figure 4.**
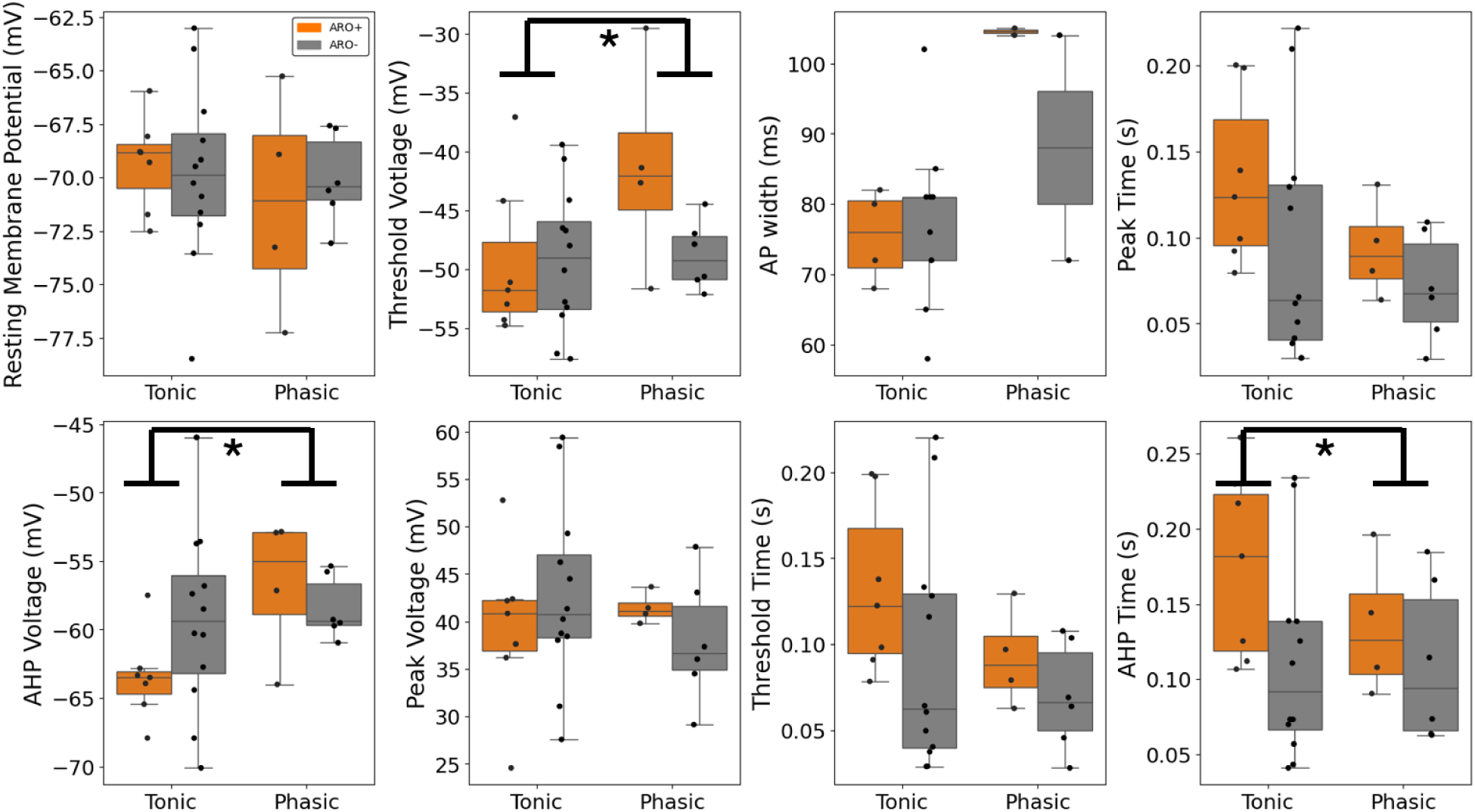
Comparison of voltage-clamp-derived PSC measurements between histologically-confirmed ARO+ (orange) and ARO- (grey) cells while considering current-clamp-derived cell phenotype (i.e. tonic vs phasic). Cell phenotype was detected as a significant factor for both Threshold Voltage and AHP Voltage, yet cell type (ARO+ vs ARO-) was not a significant factor in any of the comparisons nor were interaction significant. Each data point reflects a cell. * reflects a significant difference at an alpha of .05.

**Table 1.** Statistical summary of current-clamp recordings. For each measurement, ARO- and ARO+ cells were compared using Welch’s t-test. AHP time was found to be statistically different between ARO- and ARO+ neurons. (*) denotes significance at the .05 alpha level.

| Measurement | t-value | df (Welch) | p-value | Cohen's d | 95% CI |
| --- | --- | --- | --- | --- | --- |
| Resting Membrane Potential | -0.065 | 21.69 | 0.949 | 0.025 | [-2.86, 2.69] |
| AHP Voltage | -1.004 | 22.58 | 0.326 | 0.377 | [-6.18, 2.14] |
| Threshold Voltage | 0.937 | 14.77 | 0.364 | 0.401 | [-3.28, 8.43] |
| Voltage of Peak | -0.336 | 25.55 | 0.74 | 0.12 | [-6.93, 4.99] |
| AP width | 0.706 | 9.29 | 0.498 | 0.373 | [-11.91, 22.79] |
| Threshold Time | 1.654 | 25.08 | 0.111 | 0.597 | [-0.01, 0.07] |
| Peak Time | 1.653 | 25.09 | 0.111 | 0.597 | [-0.01, 0.07] |
| AHP time | 2.206 | 21.85 | 0.038* | 0.837 | [0.0, 0.1] |

**Table 2.** Statistical summary of current-clamp recordings considering cell phenotype (tonic or phasic) and aromatase expression (ARO- or ARO+). For each measurement, a robust trimmed factorial ANOVA was deployed to reveal main effects and the interaction between aromatase expression and cell phenotype while controlling for unequal group sizes and variances. AHP Time, AHP Voltage, and Threshold Voltage were found to be statistically different between tonic and phasic cells; no other effect or interaction was detected as significant. (*) denotes significance at the .05 alpha level.

| Measure | Effect | Q-value (value) | p-value |
| --- | --- | --- | --- |
| Peak Time | aro | 7.0516 | 0.057 |
|  | celltype | 1.0112 | 0.327 |
|  | aro × celltype | 1.2913 | 0.551 |
| AHP time | aro | 11.611 | 0.018* |
|  | celltype | 0.4075 | 0.533 |
|  | aro × celltype | 0.8642 | 0.675 |
| Threshold Time | aro | 7.0087 | 0.058 |
|  | celltype | 1.0115 | 0.327 |
|  | aro × celltype | 1.3144 | 0.545 |
| Voltage of Peak | aro | 2.1064 | 0.383 |
|  | celltype | 1.8493 | 0.188 |
|  | aro × celltype | 3.9767 | 0.176 |
| AHP Voltage | aro | 0.8263 | 0.686 |
|  | celltype | 10.3204 | 0.006* |
|  | aro × celltype | 3.7311 | 0.211 |
| Threshold Voltage | aro | 3.3496 | 0.246 |
|  | celltype | 6.9207 | 0.027* |
|  | aro × celltype | 4.7144 | 0.149 |
| Resting Membrane Potential | aro | 0.2328 | 0.9 |
|  | celltype | 0.4408 | 0.529 |
|  | aro × celltype | 0.3544 | 0.852 |

Lastly, we investigated active properties of neurons depending on their cell type and phenotype. **Figure 5** shows FI curves for phasic (**Fig. 5A**) and tonic (**Fig. 5B**) curves separately, and in terms of whether the cells were ARO+ or ARO-. Mixed ANOVA models, conducted for phasic and tonic separately, revealed no significant differences between ARO+ and ARO- cells [phasic, F(1,17) = .27, p = .61; tonic, F(1, 22) = .17, p= .68] nor any significant interactions[phasic, F(14,238) = .62, p = .85; tonic, F(14, 308) = .42, p= .97]; unsurprisingly, injected current significantly explained AP frequency for both cell types [phasic, F(14,238) = 11.08, p= 2.2e-19, 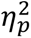= .39; tonic, F(14, 308) =25.77, p=7.4e-44, 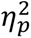= .54]. Rheobase, defined as the minimum current required to elicited a single action potential, was modeled via a 20%-trimmed mean factorial ANOVA. Cell type (Q=1.35, p=.54) and the (tonic vs phasic)X(ARO+ vs ARO-) interaction did not load significantly into the model (Q=6, p=.093). However, cell phenotype did load significantly (Q=5.07, p=.037) which suggests that, based on the direction of the data, tonic cells were more easily excitable since less current was required to elicit an action potential.

**Figure 5.**
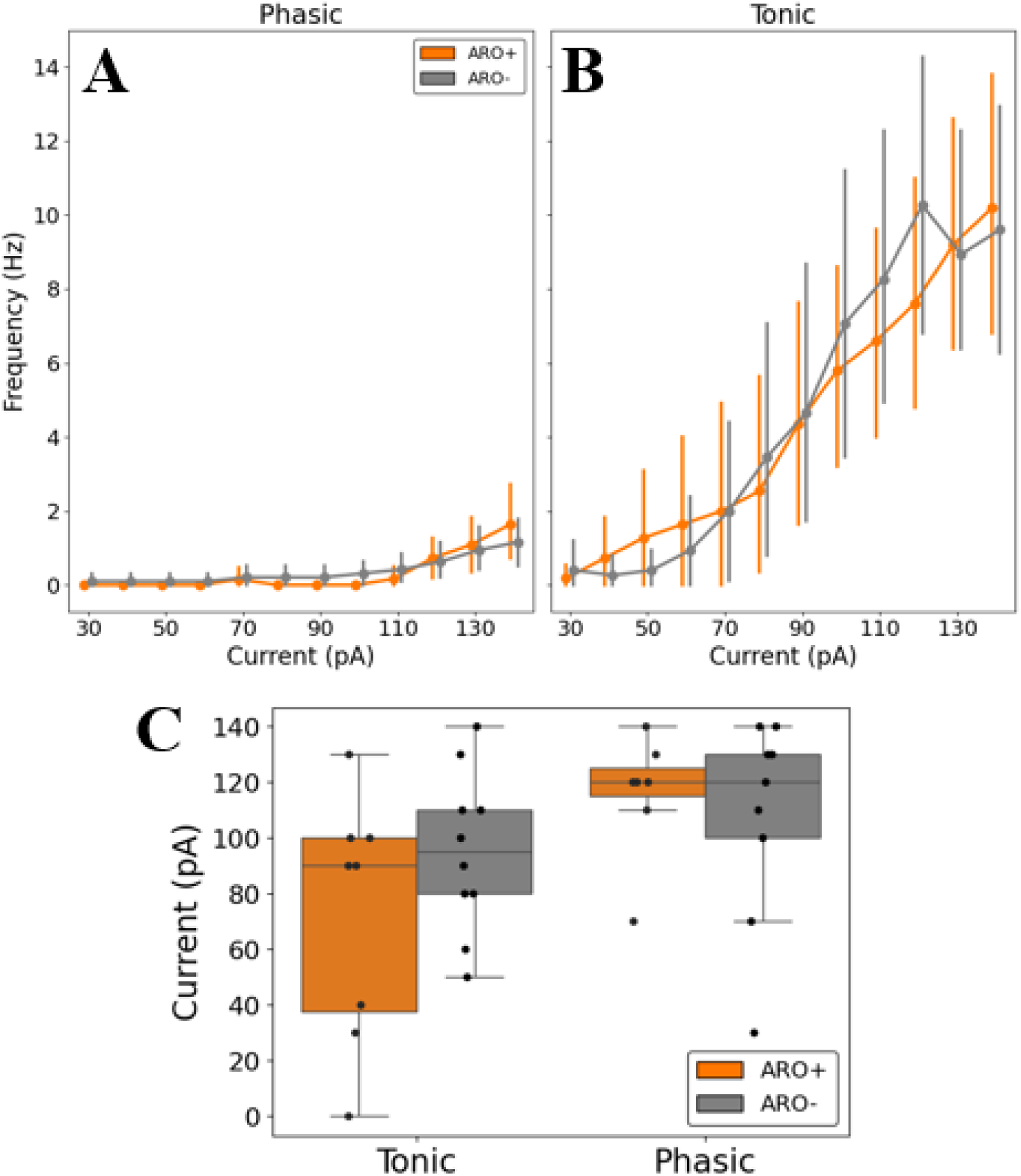
Action potential frequency as a function of current injection. The frequency of action potentials elicited by increasing current injections are displayed for phasic (A) and tonic (B), and based on aromatase expression (orange) or lack thereof (grey). C) Aromatase positivity was not a significant predictor of the current injection that elicited the first action potential (rheobase). However, cell phenotype (tonic vs phasic) significantly explained differences in rheobase, suggesting the tonic cells, with lower average rheobase values, are potentially more excitable. Each data point represents a cell

### Voltage-clamp reveals no differences between ARO+ and ARO- neurons

Voltage-clamp data were compared between cell types by measuring different metrics from PSCs: amplitude, rise time, rise slope, frequency, and slow and fast tau (**Fig. 6A**). PSC amplitude, which measures synaptic strength/conductance (e.g. the number of open ion channels) was not significantly different between ARO+ (*x̃* = 118.13, SD = 82.37) and ARO- (*x̃* = 106.84, SD = 63.39) cells, *t*(25.488) = −.716, *p* = .56, *g* = .242. Rise slope, which measures how fast synaptic conductance builds up, was found to be similar between ARO+ (*x̃* = -4.06e-9, SD = 2.66e-9) and ARO- (*x̃* = -3.34e-9, SD = 1.44e-9) cells, *t*(20.161) = .961, *p* = .494, *g* = .328. Rise time, which similarly measures build-up of synaptic conductance but is not dependent on the PSC amplitude, was also not different between ARO+ (*x̃* = 14.04, SD = .006) and ARO- (*x̃* = 13.74, SD = .006) cells, *t*(21.88) = .157, *p* = .828, *g* = .053. Lastly, the deactivation dynamics were captured by fast tau, while slow tau captured a composite of the decay kinetics of the PSC; notably, since K-gluconate internal was used, it was not possible to separate the dynamics imposed by glutamatergic or GABAergic currents, which are typically expressed as EPSCs and IPSCs. Between ARO+ (*x̃* =.011, SD = .006) and ARO- (*x̃* = .01, SD = .007) cells there was no difference in fast tau *t*(31.964) = .29, *p* = .934, *g* = .096. Slow tau was also not significantly different between ARO+ (*x̃* =.0121, SD = .006) and ARO- (*x̃* = .0123, SD = .006) cells, *t*(30.629) = −.458, *p* = .537, *g* = .152. Finally, PSC frequency, which measured the probability or density of synaptic input, was also found to not be significantly different between ARO+ (*x̃* =.79, SD = .829) and ARO- (*x̃* = .695, SD = .649) cells, *t*(27.922) = −.35, *p* = .665, *g* = .117. Median-difference 95% confidence intervals based on the Hodges-Lehmann estimator for each metric can be seen in **Figure 6B**. Together, the measured dynamics and kinetics of PSCs, using a K-gluconate internal, revealed no systematically detectable differences between ARO+ and ARO- neurons.

**Figure 6.**
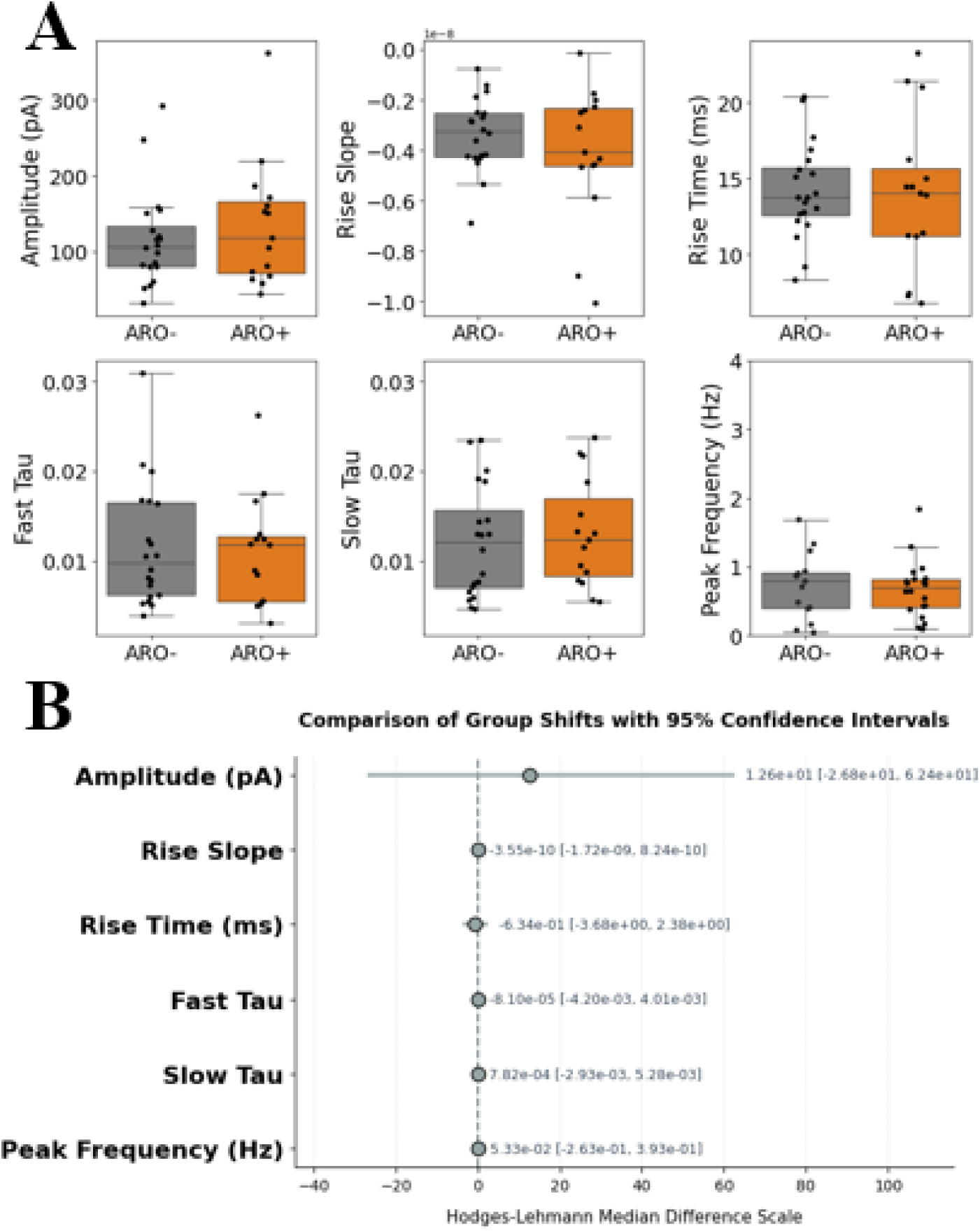
Results from voltage-clamp recordings. Within a recording, each PSC was used to measure a variety of measurements, which were collapsed for each cell separately (A). No significant differences were detected between ARO+ (orange) and ARO- (grey) cells across all computed metrics. B) 95% confidence intervals of the differences are shown for each metric. Each data point reflects a cell.

Since the above analysis relied on a pre-averaging step, such that each neuron was only represented once per analysis, we considered whether this loss of information led us to potentially overlook detectable differences on the event-by-event level. Further, our experimental cohorts included some birds of differing ages, therefore it was imperative to determine if age had an effect in our analysis. Therefore, GLMMs were used to model each metric separately, while using age as a covariate and including all PSC data (i.e. no down sampling); due to skewed distributions, all metrics were log-transformed prior to modeling via GLMM to satisfy normality assumptions. For all GLMM models, there were no detectable differences between cell types, confirming what was observed in **Figure 6**, and age was not a significant covariate suggesting that our non-significant results were not influenced by bird age.

## Discussion

Historically, estrogens have been considered as sex steroids that operate in the periphery, implicated in sexual differentiation, in part due to their circulation via the vascular system and inability to cross the blood-brain barrier. However, many reports have shown that neurons in the brain have both the capacity to synthesize neuroestrogens (via aromatase enzymatic activity) and/or to respond to estrogenic signals (via different receptor subtypes, e.g. Erα, Erβ, GPR30/GPER-1, ER-X; *for review,* Maioli et al., 2021; Toran-Allerand, 2004). Neuroestrogens are synthetized by cells that express aromatase which are found throughout the brain, including cortex. While aromatase expression is observed in cells (e.g. ovarian granulosa cells) with specialized features (e.g. low voltage-gated calcium channels; Agoston, et al., 2004) in the body, it is unclear if ARO+ neurons are themselves distinguishable from ARO- negative cells in the brain. Understanding how aromatase neurons may be distinguished by their intrinsic or network properties could help clarify the relationship between firing state and neuroestrogen synthesis, an enduring question in the field. To address this, we explored whether the intrinsic membrane properties or PSC dynamics are different between ARO+ and ARO- neurons in the songbird NCM, a region that is known for its expression of ARO+ cells, and its responsivity to, and secretion of, neuroestrogens during auditory playback.

Current-clamp recordings and analysis investigated whether intrinsic membrane properties differed between ARO+ and ARO- cells. Only a single metric, AHP Time, was found to significantly differentiate the two cell types (**Fig. 3**), all other comparisons yielded non-significant differences, which aligns well with a previous report (de Bournonville et al., 2021). This suggests that, while the two cell types are virtually indistinguishable, they differ in how the neuronal membrane potential returns to baseline following an action potential event. Specifically, a difference in AHP Time may reflect differences in potassium conductances, calcium-dependent potassium channels, and how these channels activate and deactivate. Additional analytical exploration revealed that when cell phenotype is included, there are no significant interactions, suggesting that the lack of differences between ARO+ and ARO- cells do not arise or depend on whether the cell was tonic or phasic (**Fig. 4**). Lastly, conforming with the literature, phasic cells exhibited higher rheobase—minimum current needed to trigger an action potential—relative to tonic cells (**Fig. 5C**) and tonic cells fired more action potentials than phasic cells on average (**Fig. 5A/B**). Together, the intrinsic membrane properties of ARO+ and ARO- neurons are virtually identical, suggesting no meaningful differences in ion-channel composition and concentration between the two cell types. However, it remained plausible that the inputs received and processed by these cells would make them distinguishable from each other.

Voltage-clamp electrophysiology was conducted following all current-clamp recordings on the same cells to reveal whether PSC dynamics differed between the two cell types. Out of all the measured PSC dynamics, there were no detectable differences between ARO+ and ARO-neurons. This includes the important feature of PSC frequency, which indirectly measures the extent to which the recorded cell is connected within a circuit (**Fig. 6**). All voltage-clamp recordings were performed using a K-gluconate internal solution, which introduces important constraints on the interpretation of synaptic currents. Under these conditions, the low intracellular chloride concentration shifts the chloride reversal potential near resting membrane potentials, thereby reducing the ability to resolve inhibitory currents (GABAa receptor–mediated). Consequently, inhibitory events (mIPSCs) are expected to be markedly reduced in amplitude and, in many cases, fall below detection threshold. The recorded PSCs therefore likely reflect a biased population enriched for excitatory events (mEPSCs), likely diluting the ability to separate excitatory and inhibitory contributions. In addition, because smaller-amplitude events are preferentially excluded, the analyzed PSCs may be skewed toward larger events, which can distort estimates of decay kinetics. Reduced detection of fast, low-amplitude inhibitory currents and increased noise (due to merging of unresolved inhibitory and excitatory currents) in the decay phase may lead to underestimation of fast time constants (fast tau) and poor resolution of slower components (slow tau). Accordingly, future experiments should further explore differences between ARO+ and ARO- neurons by directly assessing and comparing the dynamics of pharmacologically-restricted mIPSCs and mEPSCs separately.

Lastly, the present work explored how ARO+ and ARO− neuronal populations may differ based on their electrical and circuit properties in an *ex vivo* preparation, which precluded the ability to determine how these neurons receive chemical signals from other brain regions. For example, in NCM, dopamine-like receptors have been observed to colocalize with aromatase (Macedo-Lima et al., 2021), suggesting that this region is likely innervated by dopaminergic projections from elsewhere in the brain. This observation therefore raises the possibility that the control of, and differences between, ARO+ and ARO− neurons in NCM may arise through their interactions with distinct neurotransmitter systems originating from nearby or distant regions and, by implication, that these populations may exhibit different receptor expression profiles. Future experiments should therefore map innervation from non-NCM regions onto ARO+ and ARO− neurons within NCM, in addition to assaying the receptors they express (e.g., AMPA, NMDA, GABAa, nicotinic), to better infer their functional roles within the auditory circuit.

